# The congruence effect is impaired in healthy aging: a relationship to event-related potentials and theta-alpha oscillations

**DOI:** 10.1101/303859

**Authors:** Pau A. Packard, Tineke K. Steiger, Lluis Fuentemilla, Nico Bunzeck

## Abstract

Long-term memory can improve when incoming information is congruent with known semantic information. This so-called congruence effect has widely been shown in younger adults but age-related changes and neural mechanisms remain unclear. Here, congruence improved recognition memory in younger and older adults (i.e. congruency effect), but – importantly – this effect decreased with age. Electroencephalography data show that, in both groups, congruence led to widespread differences in event-related potentials (ERPs) and alpha-beta oscillations (8-30 Hz), known to support semantic processing. Importantly, these ERP differences predicted increases in memory performance, especially for congruent items. Finally, age-related differences in memory were accompanied by a positive ERP and later decrease in theta-alpha (5-13 Hz) during encoding, which were greater in the younger group. Together, although semantic congruence generally increases long-term memory, the effect is less pronounced in the elderly. At the neural level, theta-alpha oscillations, previously linked to memory and attentional processes, provide a mechanistic explanation for such an age-related effect.

## 1. Introduction

Long-term memory can be improved by presenting the information that has to be learned within a known context. In humans, this has often been demonstrated with a priori semantic information followed by congruent (vs. incongruent) material, and has, therefore, been labeled semantic congruence effect (or simply ‘congruence effect’) (Atienza et al., 2011; Craik and Tulving, 1975; Hall and Geis, 1980; Kapur et al., 1994; Packard et al., 2016; Schulman, 1974; Tse et al., 2011, 2007). While most previous studies have focused on younger participants (i.e. 18-35 years), the potential age-related changes and associated neural mechanisms, in particular neural oscillations, remain unclear. While age-related impairments could be expected on the basis of well-described memory deficits in the elderly, it is also clear that semantic memory (i.e. long-term memory for facts independent of time and date) is often preserved until old age (Hedden & Gabriele, 2004).

The processing of congruent semantic information per se may lead to better memories (Craik, 2002; Tibon et al., 2017). However, there is increasing support for a basic congruence dependent neural mechanism associated to the integration of memories into long-term knowledge structures or schemas (Buuren et al., 2014; Gilboa and Marlatte, 2017; Liu et al., 2017; Robin and Moscovitch, 2017; Spalding et al., 2015; Tse et al., 2011; van der Linden et al., 2017; van Kesteren et al., 2014, 2012). According to this framework, the semantic congruence matching during online encoding may be an initial step in such a general schema-dependent process of memory integration which entails interactions between the medial temporal lobe and the prefrontal cortex that favor an efficient retention and a faster consolidation of congruent events. Schema-related memory theories also nicely complement theories which emphasize the anticipatory, constructive nature of cognition and memory (Bar, 2009, 2007; Engel et al., 2001). According to this view, incoming information is linked to representations in memory that pre-activate associations, thus forming predictions and selectively facilitating cognition. Therefore, general knowledge stored in neural networks plays an important role in guiding the selection of the inputs that are meaningful according to goals.

Post-stimulus event-related potentials (ERP) during encoding reflect memory processes that are modulated by earlier input or the configuration of activity in memory (Federmeier and Laszlo, 2009; Friedman and Johnson, 2000). Specifically, semantic congruence not only drives long-term memory, as described above, but it also modulates early encoding-related post-stimulus positive ERPs (Packard et al., 2016). During healthy aging, decreases in ERPs associated with semantic congruence reflect an age-related deterioration in neural mechanisms (Huang et al., 2012; Kutas and Iragui, 1998; Wlotko et al., 2010) causing declines in encoding (Daselaar et al., 2003; Friedman et al., 1996; Glisky et al., 2001; Kamp and Zimmer, 2015; Logan et al., 2002).

In terms of neural oscillations, the beta band (16-25 Hz) has been suggested to support the maintenance of events necessary for memory encoding (Morton and Polyn, 2017). In addition, alpha-beta (8-30 Hz) oscillations may underlie encoding, possibly reflecting an increase in information (Hanslmayr et al., 2012; Hanslmayr and Staudigl, 2014), or controlled access to matching information in semantic memory (Klimesch, 2012). In general terms, theta (4-8 Hz) and alpha (8-13 Hz) oscillations closely relate to memory performance (Klimesch, 1999), and age-related changes lead to memory deficiencies (Lithfous et al., 2015). Theta-alpha oscillations may support the binding of information across large-scale networks including the prefrontal cortex and medial temporal lobe structures (Fuentemilla et al., 2014; Herweg et al., 2016). Finally, theta oscillations are involved in integrative encoding (Sans-Dublanc et al., 2017), and may mediate the positive semantic congruence effect for episodic memories and explain age-related declines (Atienza et al., 2011; Crespo-Garcia et al., 2012, 2010).

Here, we used electroencephalography (EEG) to investigate the neural mechanisms involved in the age-related differences in the semantic congruence effect. To this end, we implemented an adapted word list paradigm from a previous study (Packard et al., 2016), in which a strong memory enhancement was found for congruent words, with a corresponding early appearing ERP during encoding. Participants were presented with a series of word pairs: the first word was a semantic category (e.g. furniture) designed to preactivate specific semantic memory networks. The second word was an item either congruent (e.g. chair) or incongruent (e.g. apple) with the previous category. Subjects classified the congruence vs incongruence of the second word and their long-term recognition memory was tested in a separate subsequent phase. We expected a weaker congruence effect in the elderly as well as specific and age-related effects in post-stimulus ERPs, theta, alpha and possibly beta oscillations.

## 2. Materials and Methods

### 2.1. Participants

Thirty young (ages 19-33 years, mean 23.87; SD 3.53, 14 males) and twenty-eight elderly participants (ages 50-79 years, mean 62.55; SD 7.02, 13 males) were recruited for the behavioral experiment (experiment 1). Subsequently, twenty-three young (ages 18-28 years, mean 20.95; SD 3.23, 11 males) and twenty-five elderly participants (ages 52-79 years, mean 63.21; SD 5.82, 11 males) were recruited for the EEG experiment (experiment 2). All participants were healthy, right-handed, had normal or corrected-to-normal vision (including color-vision) and reported no history of neurological or psychiatric disorders, or current medical problems (excluding blood pressure). All elderly participants scored a 23 or higher (mean 26.45 SD 0.94) on the Montreal Cognitive Assessment version 7 (Nasreddine et al., 2005). This study and the protocol were carried out in accordance with the recommendations and approval of the local ethics committee (University of Lübeck). Each subject understood the protocol and gave written informed consent in accordance with the Declaration of Helsinki.

### 2.2. Materials

Experimental stimuli consisted of 66 categorical 6 word lists (Packard et al., 2016) translated into German, selected from category norms (Battig and Montague, 1969; Kim and Cabeza, 2007a; Yoon et al., 2004). Each list consisted of the 6 most typical instances (e.g., cow, pig, horse, chicken, sheep, and goat) of a natural/artificial category (e.g., farm animal). All of the 396 typical instances, belonging to the 66 semantic categories, were presented in separate encoding trials, each time preceded by a category (semantic cue). Additionally, semantically unrelated words were used as control words (new words) in the test phase.

### 2.3. Behavioral procedures

We used a modified version of a previous EEG experiment (Packard et al., 2017) itself adapted from other paradigms (Kim and Cabeza, 2007a, 2007b; Roediger and McDermott, 1995). Here, participants were first presented with a series of word pairs (i.e. study phase), which was followed by an informed recognition memory test. They observed the screen from a distance of 50 cm on a display with a diagonal of 62 cm. Arial letter type, 36 letter size was used.

The study phase consisted of 396 separate one word encoding trials, presented mixed in random order. Each trial started with the appearance of a fixation cross on the screen for a random duration of 2000–3000 ms. Subsequently, a category name in blue appeared on a white background for 1500 ms. After the cue disappeared, a fixation cross appeared for 2000 ms. Participants were then sequentially shown the subsequent word in green for 1000 ms. In the congruent condition, the subsequent word belonged to the semantic category (Craik and Tulving, 1975), for example, ‘colors’ followed by ‘blue’; or ‘furniture’ followed by ‘desk’. In the incongruent condition, the category name did not correspond to the subsequent word, for example ‘planets’ followed by ‘cottage’; or ‘continents’ followed by ‘oxygen’. While the second word was shown, the participants pressed a button on the mouse indicating whether the word was congruent (left click) or incongruent (right click) with the semantic category presented at the beginning of the trial. Participants were instructed to respond as quickly and correctly as possible.

Importantly, for each category, three matching words were randomly selected for the congruent condition and the remaining three words were randomly assorted with other categories for the incongruent condition. Thus, each category had an equal amount of congruent and incongruent word pairs during encoding. This controls for the possibility that differences in performance between congruent and incongruent words in the test be due to differences in participants’ memories of the category label alone, supported by guessing or generate-and-recognize strategies. There were 198 congruent-list trials and 198 incongruent-list trials. Together, the study phase lasted 50 min. At the end of this phase, subjects were presented with a distraction task in which they solved simple arithmetical problems (additions and subtractions). The distraction task ensured the participants would not rehearse the words they had previously seen. The distraction task lasted 5 min, which together with the explanations for it and the subsequent recognition test made for a total time interval of 10 min between encoding and the subsequent test. Considering that the retention interval between encoding and test was only 10 min, the paradigm was thus designed to capture only the encoding component and not the consolidation-dependent processes underlying the semantic congruence (schema) effect.

Included in the recognition test were a total of 396 Old-word (all items presented at encoding) and 396 New-word trials. The trials were presented in a pseudorandom order for each participant, thus directly avoiding any possible confounds due to order during the test. Words in the Old and New categories were predetermined and the same for all participants. Each of the 792 trials started with a fixation cross in the screen center (1500 ms). All words in the recognition phase were displayed in the middle of the screen, in green and same font and size as the study phase, each for 4000 ms. After each word, subjects responded by pressing one of 4 keys according to whether the word was judged to be “sure old,” “unsure old,” “unsure new,” or “sure new.” The scale graduations were color-coded on the keyboard. Participants were instructed to respond within 4000 ms. Every 50 trials the participants could take a short break. The test phase had a duration of 60 min approx.

### 2.4. Statistical analyses of memory results

ANOVAs (IBM SPPS Statistics 22), with encoding condition (two levels: Congruent vs Incongruent) as a within-subject factor, and age group (two levels: Young and Elderly) as a between-subject factor, were performed on the response rates and reaction times. For all analyses, *α* was set at 0.05. To estimate effect sizes, we used η_p_^2^ and Cohen’s *d* as appropriate. In the case of a participant judging a word sequence during encoding differently than we had predesigned, the sequence was not included (see Table 1). Participants’ subjective congruence ratings almost always coincided with our experimental design (94 %). Given that elderly subjects show lower memory performance for high-confidence responses (Chua et al., 2009; Dodson et al., 2007; Shing et al., 2009), we specifically ran the tests including only high-confidence responses. Corrected Hit Rates (CHR) were calculated by subtracting the proportion of erroneous ‘old’ responses from the proportion of correct ‘old’ responses. To analyze only high-confidence responses, only high-confidence correct and erroneous responses were included. Note that erroneous ‘old’ responses were for words that were not actually presented during encoding, so they were not divided into congruent or incongruent conditions. For the partial correlation analysis, sample linear correlation coefficients were calculated using the ‘partialcorr’ MATLAB function, using a Student’s *t* distribution for the transformation of the correlation for the *p*-values.

**Table 1.** Trials for EEG analyses

| Age | Trial Type | Initial Trials | TF Trials | ERP trials |
| --- | --- | --- | --- | --- |
| Young | Congruent | 180.00 | 176.52 | 144.04 |
|  | Incongruent | 186.39 | 183.17 | 142.48 |
|  | Remembered | 180.91 | 178.13 | 172.17 |
|  | Forgotten | 118.91 | 116.78 | 114.40 |
|  | Error | 29.00 |  |  |
| Elderly | Congruent | 185.04 | 181.80 | 152.20 |
|  | Incongruent | 185.28 | 181.24 | 148.04 |
|  | Remembered | 191.20 | 186.48 | 174.08 |
|  | Forgotten | 133.48 | 131.68 | 124.95 |
|  | Error | 27.48 |  |  |

Note: For both age groups the mean trials included in each category are shown. The initial trials column shows all trials before removing any trials because of EEG artifacts. Words which were incorrectly classified during encoding (e.g., congruent words classified as incongruent) are shown in the 'Error' row and were excluded from the EEG analyses. Low confidence old responses were not included in the 'Remembered' category. Mean trials included in the TF analysis are shown in the 'TF Trials' column, and trials included in the ERP analysis are shown under 'ERP Trials'.

### 2.5. EEG analysis – experiment 2

During encoding, electroencephalographic (EEG) activity was acquired with an Easy Cap system by BrainProducts with 32 standard active electrodes. For detecting vertical and horizontal eye movement (VEOG / HEOG), 4 electrodes were used. Impedances were maintained under 20 kΩ. FCz served as reference and AFz as ground electrode. The sampling rate was at 500 Hz with online high-pass (0.1 Hz) and low-pass (240 Hz) filters. EEGLAB (version 13; Delorme and Makeig, 2004) and customized MATLAB version 2016b (The MathWorks) tools were used for preprocessing the EEG data offline.

### 2.6. ERP analysis

For the ERP analysis, a filter appropriate for slow components was used (Acunzo et al., 2012; Tanner et al., 2016). Specifically, the data were filtered offline with a low-pass filter set at 60 Hz, with the default ‘pop_eegfiltnew’ EEGLAB function, with no additional high-pass filtering as the online high-pass filter (0.1 Hz) was deemed sufficient. Secondly, all trials of the encoding phase were epoched and down sampled to 250 Hz. ERPs during the encoding were studied by extracting event-locked EEG epochs of 1600 ms, ending 1500 ms after stimulus onset, with the 100 ms prior to stimulus onset used for the baseline. Subsequently, major atypical artifacts, trials with amplifier saturation, and bad channels were visually identified and removed, (maximum 5 channels, mean = 1.2). Afterwards, blinks and eye movement artifacts were removed with independent component analysis (ICA; Delorme and Makeig, 2004)). Finally, bad channels were interpolated. Oz was selected to re-reference the data, as re-referencing to average can mask the effects of EEG differences with a broad distribution across the scalp (Luck, 2005), such as we expected following previous experiments (Packard et al., 2016). EEG trials with a shift exceeding 100 μV were rejected offline.

For the congruence analysis, only the trials correctly classified during encoding were divided into two separate conditions, ‘Congruent’, and ‘Incongruent’ (including both subsequently remembered and forgotten trials, see Table 1 for mean amount of trials for each condition, including mistakes). For the subsequent memory analysis, only trials correctly classified during encoding were divided into two separate conditions, ‘Remembered’, and ‘Forgotten’ (including both congruent and incongruent trials). To increase the reliability of the subsequent memory contrast, only high-confidence correct ‘old’ responses were included in the ‘Remembered’ condition. High and low confidence misses (‘new’ responses for words actually presented during the encoding phase) were included in the ‘Forgotten’ condition. Three young subjects and one elderly subject were excluded from the analysis due to excessively noisy data or being a low performing outlier (with exceedingly low HR compared to their age group as identified with SPSS using a step of 1.5 x Interquartile Range). Fieldtrip (Oostenveld et al., 2010) and customized MATLAB scripts were used for statistical data analysis.

To detect reliable differences between the conditions during encoding, the conditions were contrasted using Fieldtrip via a two-tailed non-parametric cluster-based permutation test (Maris and Oostenveld, 2007). All time points between 50 ms and 1500 ms at 27 scalp electrodes (Fp1, Fp2, F7, F3, Fz, F4, F8, FC5, FC1, FC2, FC6, T7, C3, Cz, C4, T8, CP5, CP1, CP2, CP6, P7, P3, Pz, P4, P8, O1, O2) were included in the test. Time points before 50 ms were too early to be considered relevant for the test. For all contrasts, a *t* test was performed for each sample (channel, time). For each permutation, all t scores corresponding to uncorrected *p* values of 0.05 were formed into clusters. The sum of the t scores in each cluster is the ‘mass’ of that cluster and the most extreme cluster mass in each of the sets of tests was recorded and used to estimate the distribution of the null hypothesis. The Monte Carlo estimate was calculated by running random permutations of the condition labels (*n* = 1000) and comparing the cluster statistics found in the real data with that found in the random data. The p-value is thus obtained with the proportion of cluster statistics in the random data exceeding that in the real data. Clusters were formed from significant samples (*p* < 0.05), considering only effects with minimum three significant neighboring channels based on triangulation.

### 2.7. TF analysis

For the TF analysis, the data were high-pass (0.5 Hz) and low-pass (35 Hz) filtered. Second, all trials of the encoding phase were epoched and down sampled. We extracted event-locked EEG epochs of 4500 ms starting at 2000 ms before the presentation of the first word of the word list. Subsequently, major atypical artifacts, trials with amplifier saturation, and bad channels were visually identified and removed (maximum 1 channel per subject). Otherwise, the preprocessing was conducted as described above for the ERPs.

TF decompositions were conducted from 2 Hz to 30 Hz, from 1000 ms before stimulus onset to 1500 ms after stimulus onset, using convolution on the single-trial time series with complex Morlet wavelets (4 cycles), with steps of 8 ms in the time and 0.22 Hz in the frequency domain. For each condition, power was averaged across trials. A 300 ms baseline correction was applied (from 500 ms before stimulus onset to 200 ms before stimulus onset). The power values thus obtained indicated the relative change as compared to the power during the baseline period, that is, a value of 1 would indicate no change respect to baseline. Note that in the statistical tests and the figures, we subtracted different conditions from each other, so that a value of 0 in those contrasts would indicate no differences between conditions.

To detect reliable differences between the conditions during encoding, the conditions were contrasted using Fieldtrip via a two-tailed non-parametric cluster-based permutation test (Maris and Oostenveld, 2007), on the frequency range from 2 Hz to 30 Hz. For all contrasts, a *t* test was performed for each sample (channel, frequency, time), the rest of the analysis was performed as described above for the ERPs.

## 3. Results

### 3.1. Behavioral findings

#### 3.1.1. Experiment 1

##### 3.1.1.1. Main effects of congruence, and age, and a congruence by age interaction, for high-confidence corrected hit rate

The proportions of high-confidence ‘Sure’ responses during the recognition phase were analyzed (see Figure 1A, and Table 2 for hit and false-alarm rates including low and high confidence responses). A 2×2 ANOVA with the factors congruence and age revealed a significant main effect of congruence (*F*_(1,56)_ = 515.45, *p* < 0.001, ηp^2^ = 0.90), driven by higher CHR for congruent words (mean 0.64, SEM 0.02), than for incongruent words (mean 0.41, SEM 0.02). There was also a significant main effect of age (*F*_(1,56)_ = 4.72, *p* = 0.034, ηp^2^ = 0.08), with higher CHR for younger (mean 0.60, SEM 0.02), than for elderly subjects (mean 0.53, SEM 0.03). Importantly, a significant congruence by age interaction effect was also revealed (*F*_(1,56)_ = 4.73, *p* = 0.034, ηp^2^ = 0.08). Post-hoc t-tests showed that young subjects had higher congruent corrected hit-rates (*t*(56) = 2.63, *p* = 0.011, Cohen’s d = 0.69). In contrast, there were no significant differences between incongruent hit rates between young and elderly subjects (*p* = .133).

**Figure 1.**
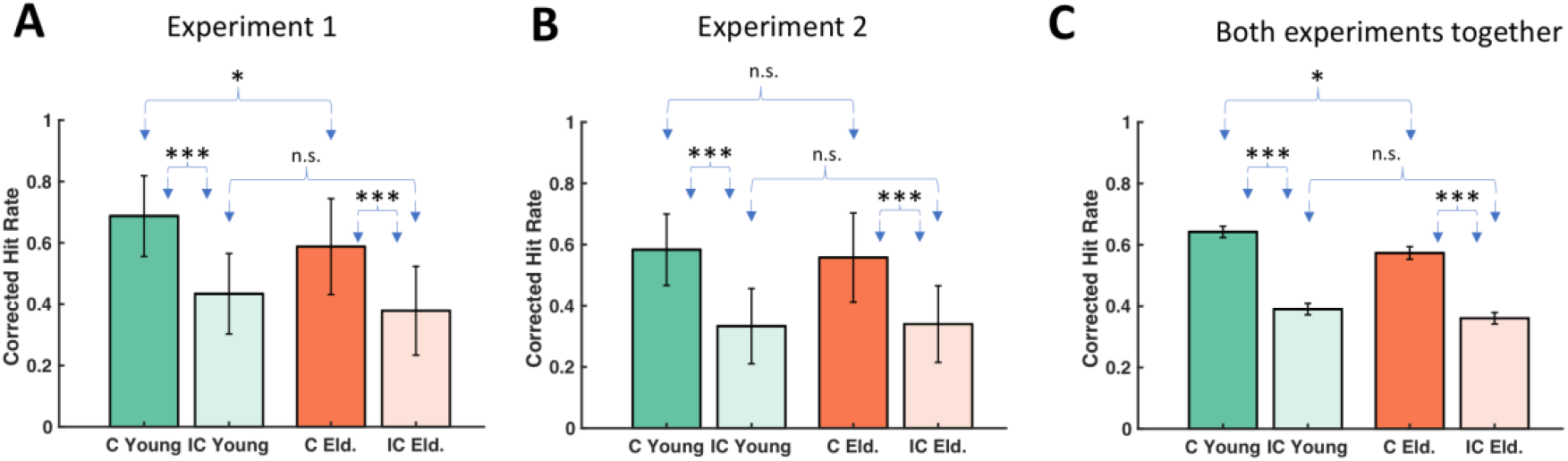
(A) CHR for Experiment 1. All conditions were above chance level. There was a main effect of congruence, a main effect of age, and a congruence by age interaction. (B) CHR for Experiment 2. All conditions were above chance level. There was a main effect of congruence, but the effect of age and the congruence by age interaction were not significant. (C) CHR for both experiments together. All conditions were above chance level. There was a main effect of congruence, and a congruence by age interaction. The effect of age was not significant. \**p* < 0.05. *** *p* < 0.001. Error bars indicate SEM.

**Table 2.** Memory performance

| Group | Category | All Responses | Sure Responses |
| --- | --- | --- | --- |
| Experiment 1 |  |  |  |
| Young | Congruent HR | .84 (0.02) | .72 (.02) |
|  | Incongruent HR | .64 (0.02) | .47 (.03) |
|  | False Alarm rate | .09 (.02) | .04 (.01) |
| Elderly | Congruent HR | .75 (.03) | .64 (.03) |
|  | Incongruent HR | .56 (.03) | .43 (.03) |
|  | False Alarm rate | .09 (.02) | .05 (.01) |
| Experiment 2 |  |  |  |
| Young | Congruent HR | .77 (.02) | .62 (.03) |
|  | Incongruent HR | .57 (.02) | .37 (.03) |
|  | False Alarm rate | .09 (.02) | .04 (.01) |
| Elderly | Congruent HR | .74 (.03) | .63 (.03) |
|  | Incongruent HR | .54 (.03) | .41 (.04) |
|  | False Alarm rate | .11 (.02) | .07 (.02) |

Note: Mean and SEM for Hit Rates (HR) and False-Alarm rates for both experiments and age groups are shown. Under the 'All Responses' column both high confidence and low confidence 'Old' responses are shown. Under the 'Sure Responses' column only high confidence responses are included.

##### 3.1.1.2. Main effect of congruence for RT

For participants’ reaction times during the encoding phase, we found a main effect of congruence (*F*_(1,56)_ = 44.04, *p* < 0.001, ηp^2^ = 0.44), participants were slower at identifying incongruent words (see Table 3). However, age did not reach significance level (*F*_(1,56)_ = 2.29, *p* = 0.136, ηp^2^ = 0.04), and there was no significant congruence by age interaction (*F*_(1,56)_ = 0.62, *p* = 0.435, ηp^2^ = 0.01).

**Table 3.**
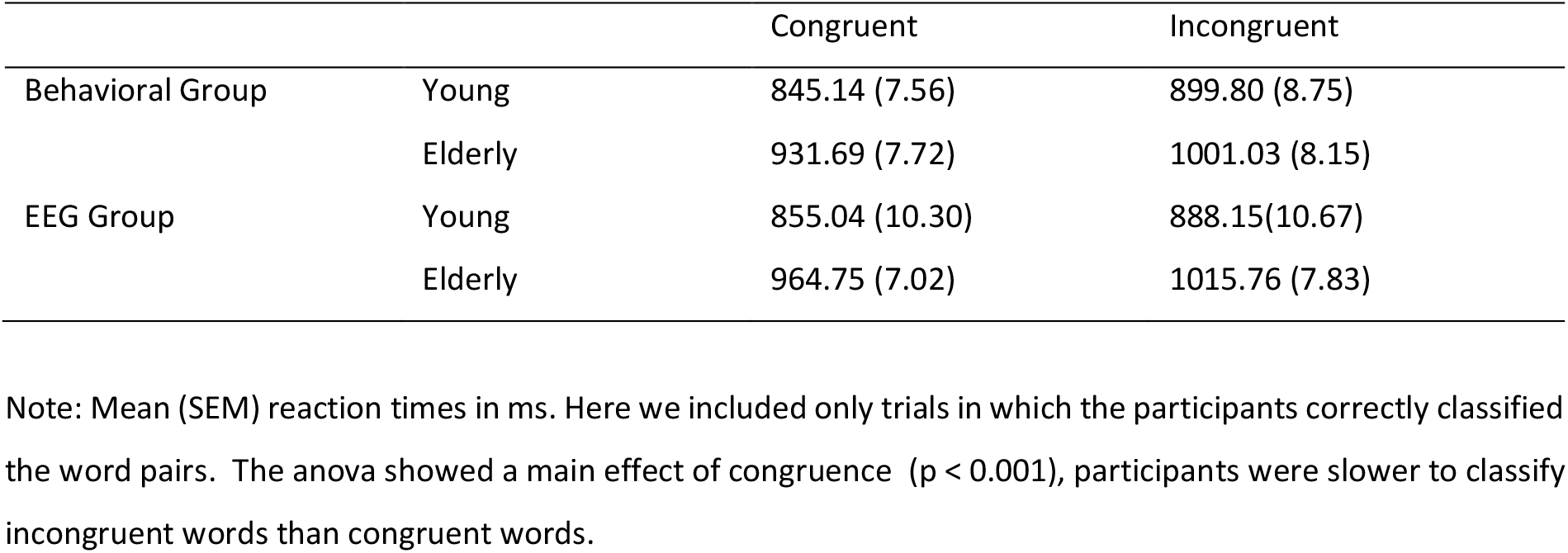
Reaction times for the semantic matching encoding task

|  |  | Congruent | Incongruent |
| --- | --- | --- | --- |
| Behavioral Group | Young | 845.14 (7.56) | 899.80 (8.75) |
|  | Elderly | 931.69 (7.72) | 1001.03 (8.15) |
| EEG Group | Young | 855.04 (10.30) | 888.15(10.67) |
|  | Elderly | 964.75 (7.02) | 1015.76 (7.83) |

Note: Mean (SEM) reaction times in ms. Here we included only trials in which the participants correctly classified the word pairs. The anova showed a main effect of congruence ( $p < 0.001$ ), participants were slower to classify incongruent words than congruent words.

#### 3.1.2. Experiment 2

##### 3.1.2.1. Main effect of congruence for high-confidence CHR

In experiment 2, the proportions of high-confidence ‘Sure’ responses during the recognition phase were analyzed, again, in a 2×2 ANOVA (see Figure 1B, and Table 2 for hit and false-alarm rates including low and high confidence responses). This analysis only showed a significant main effect for congruence (*F*(1,46) = 292.01, *p* < 0.001, ηp^2^ = 0.86) with higher CHR for congruent words (mean 0.57, SEM 0.02), than for incongruent words (mean 0.34, SEM 0.03). In contrast to the behavioral experiment (Experiment 1), there was no main effect of age (*F*(1,46) = 0.075, *p* = 0.785, ηp^2^ < 0.01), and no congruence by age interaction effect (*F*(1,46) = 1.39, *p* = 0.244, ηp^2^ = 0.03).

##### 3.1.2.2. Main effect of congruence for RT

For participants’ reaction times during the encoding phase, there was a main effect of congruence (*F*(1,46) = 22.42, *p* < 0.001, ηp^2^ = 0.33), participants were slower at identifying incongruent words (see Table 3). However, age did not quite reach significance level (*F*(1,46) = 3.59, *p* = 0.06, ηp^2^ = 0.07), and there was no significant congruence by age interaction (*F*(1,46) = 1.02, *p* = 0.319, ηp^2^ = 0.02). This result on reaction times replicated the results of the behavioral experiment and is in line with the recognition memory performance (i.e. no age effect and no interaction).

#### 3.1.3. Analysis of both experiments

##### 3.1.3.1. Main effect of congruence, experiment group, and a congruence by age interaction, for high-confidence CHR

A separate 2×2×2 ANOVA on high-confidence ‘Sure’ responses during the recognition phase, with congruence, age group, and experimental group as factors, including all the participants from both experiments, revealed a significant main effect of congruence (*F*(1,102) = 772.28, *p* < 0.001, ηp^2^ = 0.88); a marginal effect of age (*F*(1,102) = 3.00, *p* = 0.087, ηp^2^ = 0.029); and a main effect of experiment (*F*(1,102) = 7.39, *p* = 0.008, ηp^2^ = 0.068), with lower CHR in the EEG group (see Figure 1C). Importantly, there was a significant congruence by age interaction effect (*F*(1,102) = 5.24, *p* = 0.024, ηp^2^ < 0.05). This interaction was driven by a stronger congruence effect in the younger subjects as compared to the elderly. Specifically, post-hoc t-tests showed the young subjects had higher congruent high-confidence corrected hit-rates (*t*(104) = 2.47, *p* = 0.015, Cohen’s *d* = 0.48). In contrast, there were no significant differences between incongruent high-confidence hit rates between young and elderly subjects (*p* = .261). Finally, there were no other interactions between experimental group and age or congruence (all *p* > 0.170).

##### 3.1.3.2. Main effect of congruence, and age, for RT

There was a main effect of congruence for participants’ reaction times (*F*_(1,102)_ = 63.37, *p* < 0.001, ηp^2^ = 0.38), participants were slower at identifying incongruent words (see Table 3). There was also a significant main effect of age (*F*_(1,102)_ = 5.72, *p* = 0.019, ηp^2^ = 0.05), younger subjects were faster, but there was no significant congruence by age interaction (*F*_(1,102)_ = 1.55, *p* = 0.216, ηp^2^ = 0.02). There was no significant main effect of EEG group (*F*_(1,102)_ = 0.67, *p* = 0.796, ηp^2^ = 0.1), and no other interactions between EEG group and age or congruence (all *p* > 0.129).

### 3.2. EEG findings

#### 3.2.1. ERP Cluster analysis

##### 3.2.1.1. Main effect of congruence, and a congruence by age interaction

A Monte Carlo cluster-based permutation test was performed on the data of young and elderly grouped together, from 50 ms to 1500 ms after word onset, which revealed a positive cluster due to congruence during encoding, from 320 ms to 1176 ms, with a broad central topography (congruent vs incongruent, see Table 4). Next, the effect of congruence between young and elderly was contrasted in the same way (congruent minus incongruent in young subjects vs congruent minus incongruent in elderly subjects), and significant differences in a positive cluster were found, from 404 ms to 604 ms, with a broad central topography (see Figure 2). As such, we found a main effect of congruence and an interaction between age and congruency that resembles the behavioral effects.

**Figure 2.**
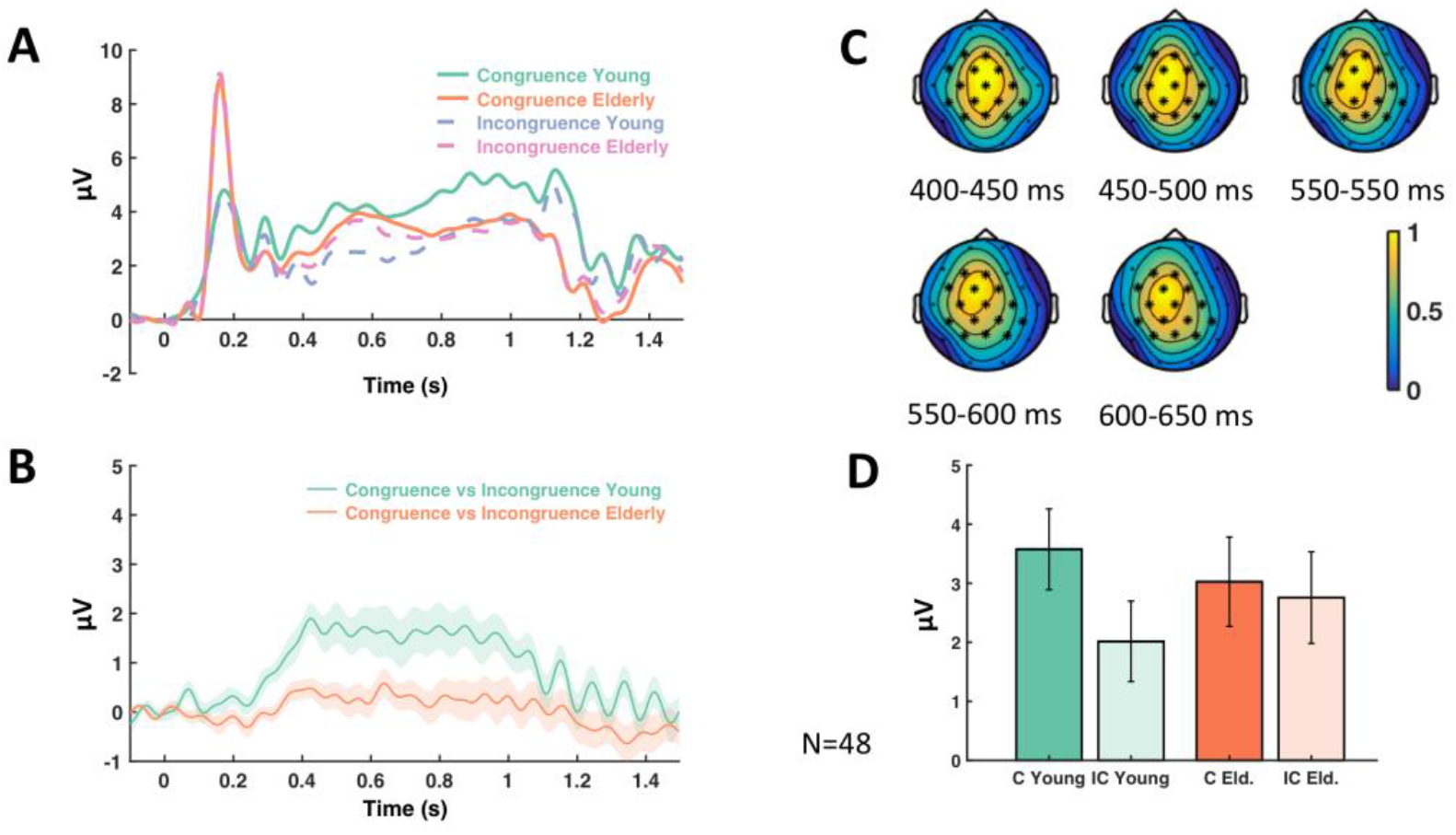
ERP congruence effect. (A) Mean amplitude of the congruence by age ERP interaction across the significant electrodes found in the cluster. The time window of the significant differences detected in the cluster was from 404 to 604 ms. (B) The difference wave of the congruent minus the incongruent condition in the young group was contrasted against the same difference wave in the elderly group, in the cluster analysis. For the figure, the mean was calculated from the grand averages across the 15 electrodes (F3, Fz, FC5, FC1, FC2, C3, Cz, C4, CP5, CP1, CP2, CP6, P3, Pz, P4) of the cluster. (C) ERP Topoplots (mean amplitude in *μV*) of the significant cluster of the congruence by age interaction. Electrodes where significant differences were detected corresponding to the interaction are highlighted with asterisks. The difference wave of the congruent minus the incongruent condition in the young group was contrasted with the same difference wave in the elderly group in the cluster analysis. (D) Barplot showing the mean amplitude across the electrodes of the significant cluster of the congruence by age interaction, from 404 to 604 ms, with the congruent (c) and incongruent condition (ic) shown for the groups of young and elderly (eld.) adults.

**Table 4.** Significant Clusters

Significant Clusters
| Cluster | Duration | Frequency range | Electrodes | p |
| --- | --- | --- | --- | --- |
| Congruence contrast: positive ERP | 320-1176 ms | - | Fz F4 FC1 FC2 FC6 C3 Cz C4 T8 CP1 CP2 CP6 Pz P4 | 0.002 |
| Congruence by age interaction: positive ERP | 404-604 ms | - | F3 Fz FC5 FC1 FC2 C3 Cz C4 CP5 CP1 CP2 CP6 P3 Pz P4 | 0.012 |
| Subsequent Memory contrast: positive ERP | 224-1496 ms | - | F3 Fz F4 FC5 FC1 FC2 FC6 T7 C3 Cz C4 T8 CP5 CP1 CP2 CP6 P3 Pz P4 | 0.002 |
| Subsequent Memory by age interaction: positive ERP clusters 1-3 | 408-612 ms;<br>620-684 ms;<br>724-1172 ms | - | F3 Fz F4 FC5 FC1 FC2 FC6 (T7)* C3 Cz C4 T8 CP5 CP1 CP2 CP6 P3 Pz P4 | 0.002;<br>0.022;<br>0.002; |
| Congruence contrast: relative power decrease (TF) | 50-1440 ms | 2-25.56 Hz | F3 Fz F4 FC5 FC1 FC2 FC6 T7 C3 Cz C4 T8 CP5 CP1 CP2 CP6 P3 Pz P4 | 0.002 |
| Congruence by age interaction: relative power decrease (TF) | 728-1072 ms | 4.89-13.56 Hz | F3 Fz F4 FC5 FC1 FC2 FC6 T7 C3 Cz C4 T8 CP5 CP1 CP2 CP6 P3 Pz P4 | 0.030 |
| Remembered minus forgotten (TF) | 600-1336 ms | 2.67-28.44 Hz | F3 Fz F4 FC5 FC1 FC2 FC6 T7 C3 Cz C4 T8 CP5 CP1 CP2 CP6 P3 Pz P4 | 0.002 |

Note: Contrasts for each cluster are indicated as detected by the two-tailed non-parametric cluster-based permutation tests. ERP and TF clusters are presented. Congruence contrasts (congruent minus incongruent), congruence by age interaction contrasts (Congruence contrast in young group minus congruence contrast in elderly group), subsequent memory contrasts (remembered minus forgotten) and subsequent memory by age interaction contrasts (subsequent memory contrast in young group minus subsequent memory contrast in elderly group) are shown. The duration in ms and the list of electrodes which were detected in the cluster are shown for each contrast, together with the significance level. \* T7 is only present in subsequent memory by age interaction: positive ERP clusters 1 and 3.

##### 3.2.1.2. Main effect of subsequent memory, and a subsequent memory by age interaction

The cluster-based permutation test, on the data of young and elderly grouped together, from 50 ms to 1500 ms after word onset, revealed a significant positive cluster of Differences due to Memory, from 224 ms to 1496 ms, with a broad central topography (DM effect; remembered vs forgotten, see Table 4). When the DM effect between young and elderly was contrasted in the same way (remembered minus forgotten in young subjects vs remembered minus forgotten in elderly subjects), significant differences in three different positive clusters were found similar to the congruence effects, the first cluster starting at 408 ms and the last ending at 1172 ms, all with a broad central topography (see Figure 3). Note that a DM analysis could not be calculated for congruent vs. incongruent words due to a lack of sufficient numbers of trials per condition after artifact rejection. Specifically, the remembered congruent category was much lower than the others (see Table 5), especially for some of the subjects.

**Figure 3.**
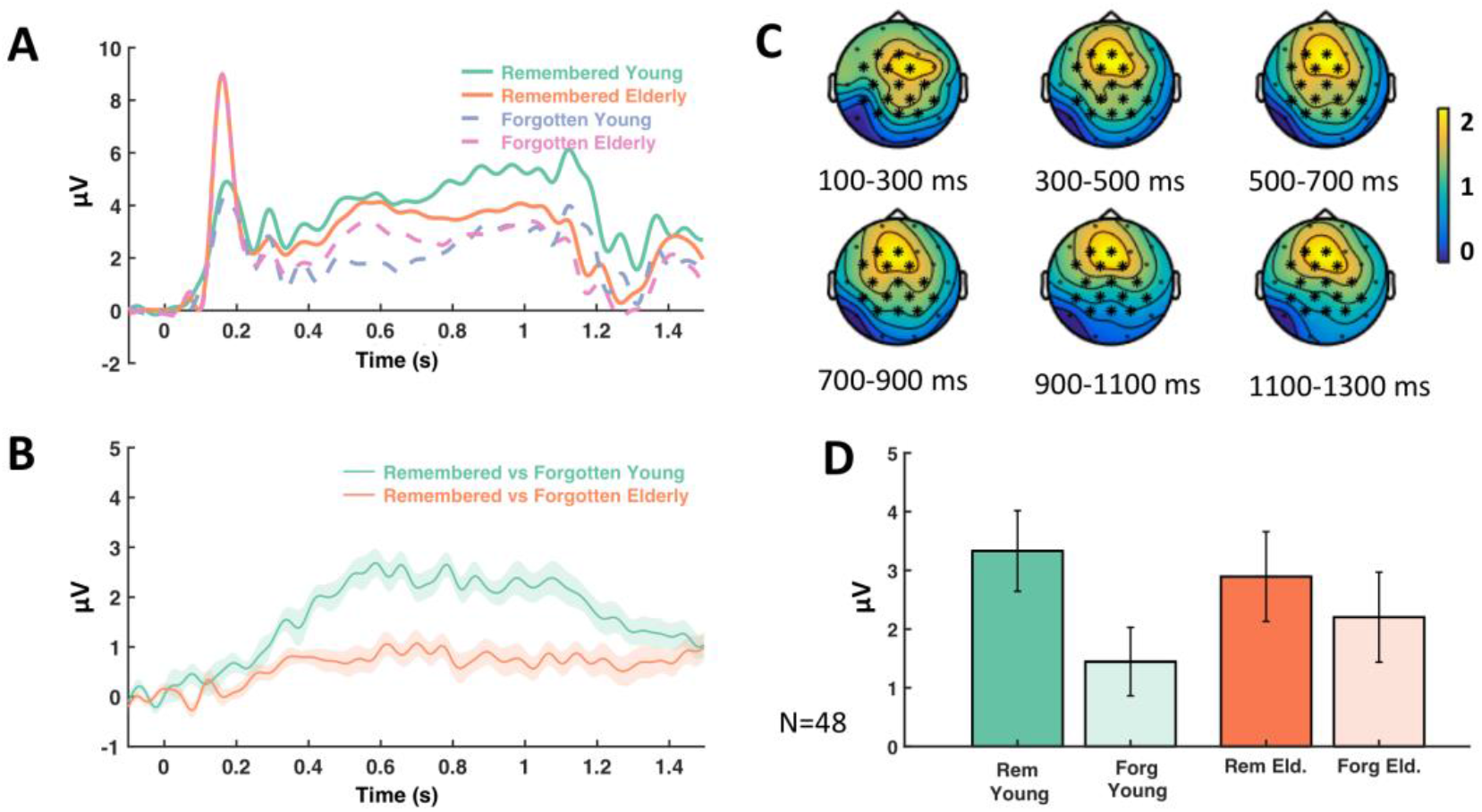
Memory by Age ERP interaction. (A) Mean amplitude of the memory by age ERP interaction across the significant electrodes found in the cluster. The time windows of the significant differences detected in the three clusters were from 408 to 612 ms, 620 to 684 ms, and 724 to 1172 ms. (B) The difference wave of the remembered minus the forgotten condition in the young group was contrasted against the same difference wave in the elderly group, in the cluster analysis. For the figure, the mean was calculated from the grand averages across the 15 electrodes (F3, Fz, FC5, FC1, FC2, C3, Cz, C4, CP5, CP1, CP2, CP6, P3, Pz, P4) of the cluster. (C) ERP Topoplots (mean amplitude in *μV*) of the significant cluster of the memory by age interaction. Electrodes where significant differences were detected corresponding to the interaction are highlighted with asterisks. The difference wave of the congruent minus the incongruent condition in the young group was contrasted with the same difference wave in the elderly group in the cluster analysis. (D) Barplot showing the mean amplitude across the electrodes of the significant cluster of the memory by age interaction, from 408 to 1172 ms, with the remembered (rem) and forgotten condition (forg) shown for the groups of young and elderly (eld.) adults.

**Table 5.**

|  | Congruent<br>Remembered | Incongruent<br>Remembered | Congruent<br>Forgotten | Incongruent<br>Forgotten |
| --- | --- | --- | --- | --- |
| Young | 105 | 64.13 | 39.70 | 77.43 |
| Elderly | 100.38 | 60.92 | 42.58 | 81.92 |

Note: For both age groups the mean trials for the congruent remembered, incongruent remembered, congruent forgotten and incongruent forgotten subcategories are shown. Only clean trials as remained after artifact rejection are shown.

#### 3.2.2. Time-Frequency Cluster Analysis

##### 3.2.2.1. Main effect of congruence, and a congruence by age interaction

A Monte Carlo cluster-based permutation test was run on the TF data of young and elderly grouped together (see Table 4), from 50 ms to 1500 ms after word onset, and from 2 to 30 Hz. It revealed significant differences due to congruence (congruent vs incongruent) in a negative cluster, with a broad central topography, approximately from 2 Hz to 26 Hz, from 50ms to 1440 ms (see Figure 4). In a next step, the differential effect of congruence between young and elderly was contrasted in the same way (congruent minus incongruent in young subjects vs congruent minus incongruent in elderly subjects). This analysis revealed significant differences (see Figure 5) in a negative cluster, with a broad central topography, approximately from 5 Hz to 14 Hz, from 728 ms to 1072 ms after stimulus onset. Again, this activity pattern (i.e., a main effect of congruence and interaction between age and congruency) mirrors the behavioral effect.

**Figure 4.**
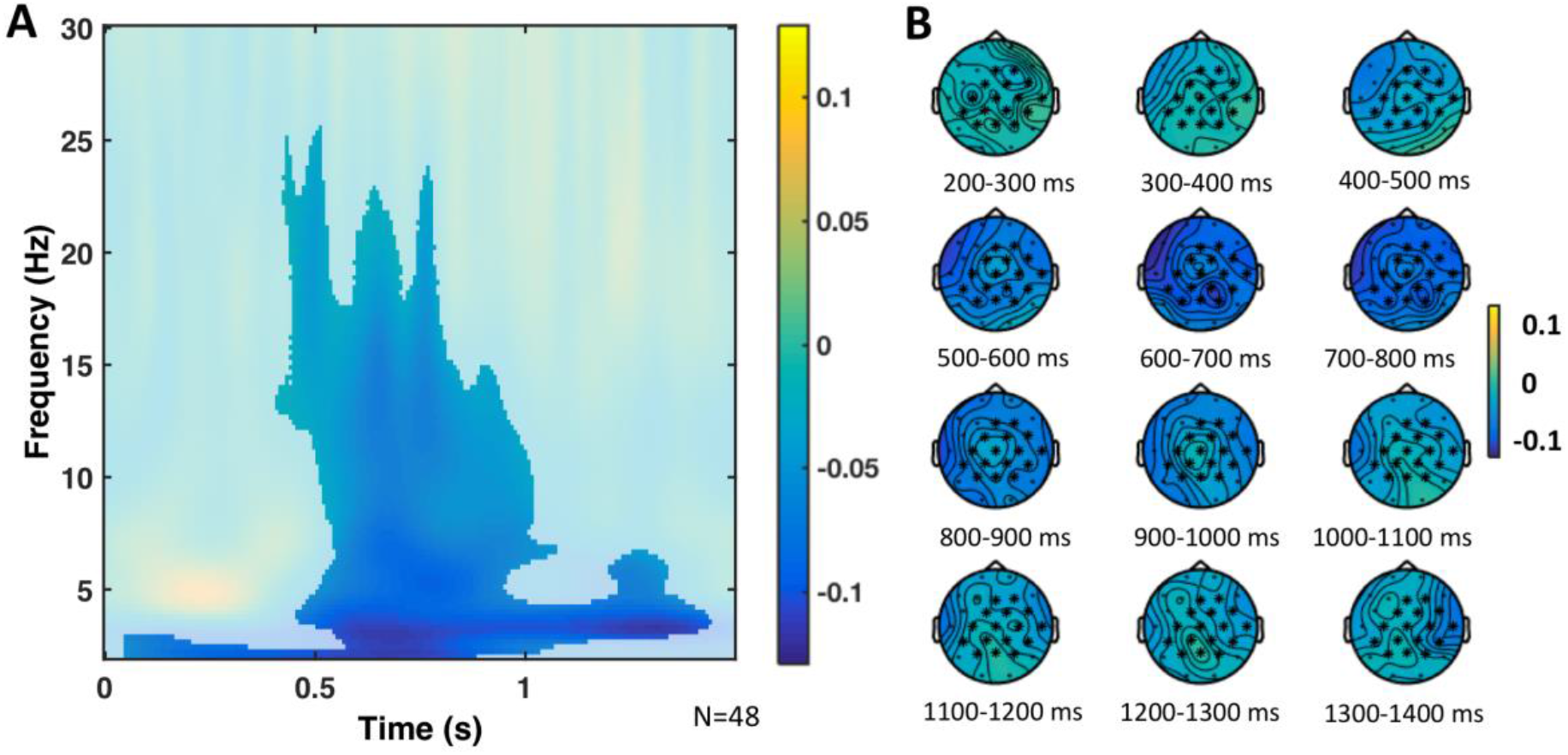
Congruence TF effect. (A) Relative power change of the congruence TF effect across the significant electrodes found in the cluster. The significant time window was from 50 ms to 1440 ms, and the significant frequency window was from 2 Hz to 25.56 Hz. The difference between the congruent minus the incongruent condition was contrasted in the cluster analysis, including both the young and elderly groups. For the figure, the mean was calculated from the grand averages across the 19 electrodes (F3, Fz, F4, FC5, FC1, FC2, FC6, T7, C3, Cz, C4, T8, CP5, CP1, CP2, CP6, P3, Pz, P4) of the cluster. (B) TF power topoplots of the significant cluster of the congruence effect, in the significant TF window. Electrodes in the cluster are highlighted with asterisks.

**Figure 5.**
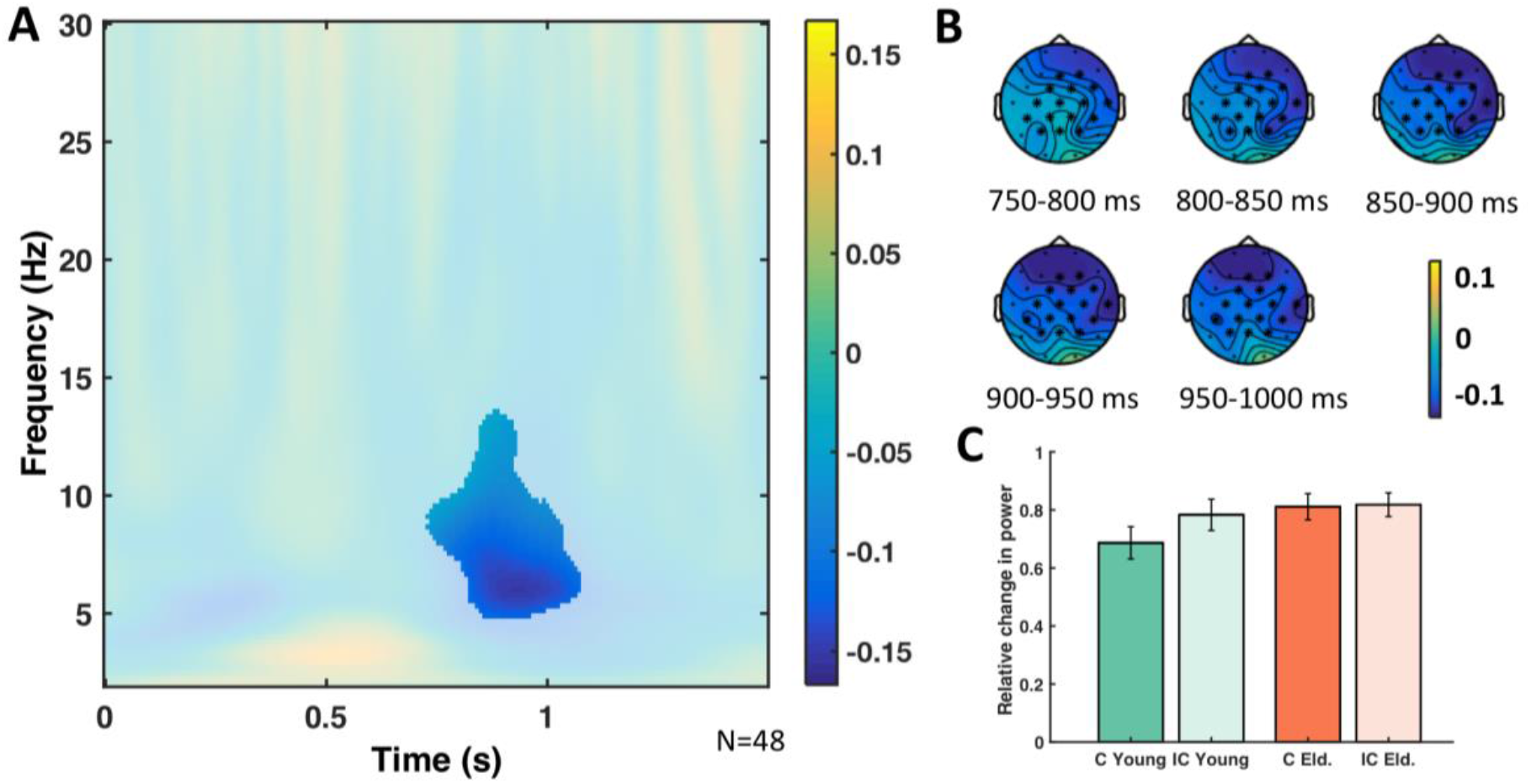
Congruence by Age TF interaction. (A) Relative power change of the congruence by age TF interaction across the significant electrodes found in the cluster. The significant time window was from 728 ms to 1072 ms, and the significant frequency window was from 4.89 Hz to 13.56 Hz. The difference in relative power of the congruent minus the incongruent condition in the young group was contrasted against the same difference in the elderly group, in the cluster analysis. For the figure, the mean was calculated from the grand averages across the 19 electrodes (F3, Fz, F4, FC5, FC1, FC2, FC6, T7, C3, Cz, C4, T8, CP5, CP1, CP2, CP6, P3, Pz, P4) of the cluster. (B) TF power topoplots of the significant cluster of the congruence by age interaction, in the significant TF window. Electrodes with significant differences corresponding to the interactions are highlighted with asterisks. (C) Barplot showing the relative power change across the electrodes of the significant cluster of the congruence by age interaction, in the significant TF window, with the congruent (c) and incongruent condition (ic) shown for the groups of young and elderly (eld.) adults.

##### 3.2.2.2. Main effect of subsequent memory, and a congruence by age interaction

The cluster-based permutation test, on the data of young and elderly grouped together, from 50 ms to 1500 ms after word onset, and from 2 to 30 Hz, revealed significant differences in a negative cluster due to memory (remembered vs forgotten, see Figure 6, and Table 4), from 600 ms to 1336 ms, from approximately 3 Hz to 28 Hz. When we contrasted for differences due to memory between young and elderly in the same way (remembered minus forgotten in young subjects vs remembered minus forgotten in elderly subjects), we did not find any significant differences. Note that a DM analysis could not be calculated for congruent vs. incongruent words due to a lack of sufficient numbers of trials per condition (see Table 5), as explained above.

**Figure 6.**
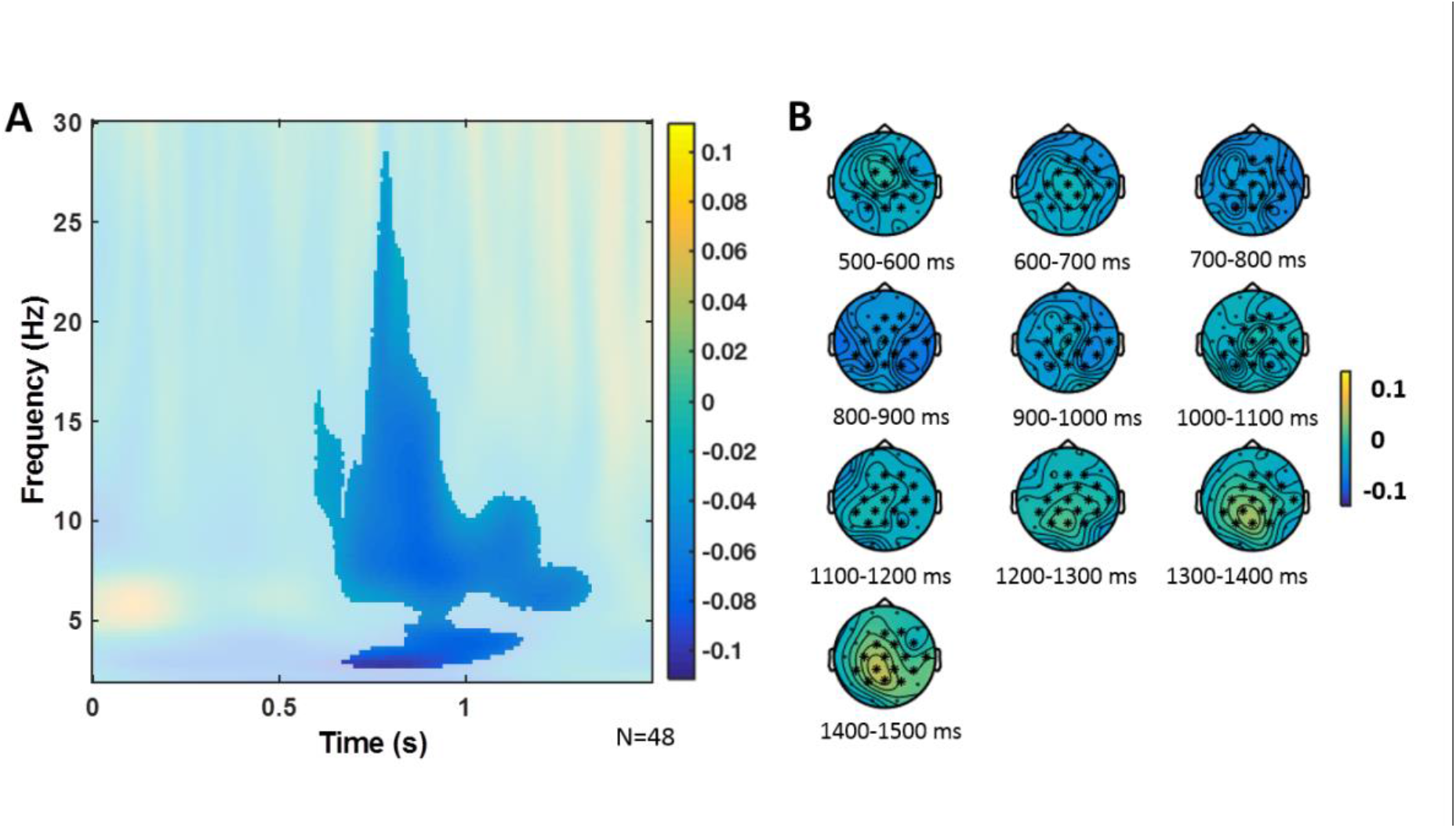
Subsequent Memory TF effect. (A) Relative power change of the subsequent memory TF effect across the significant electrodes found in the cluster. The significant time window was from 600 ms to 1336 ms, and the significant frequency window was from 2.66 Hz to 28.44 Hz. The difference between the remembered minus the forgotten condition was contrasted in the cluster analysis, including both the young and elderly groups. For the figure, the mean was calculated from the grand averages across the 19 electrodes (F3, Fz, F4, FC5, FC1, FC2, FC6, T7, C3, Cz, C4, T8, CP5, CP1, CP2, CP6, P3, Pz, P4) of the cluster. (B) TF power topoplots of the significant cluster of the subsequent memory effect, in the significant TF window. Electrodes in the cluster are highlighted with asterisks.

#### 3.2.3. Behavioral EEG correlations

To investigate whether the congruence effect directly relates to neural activity, a partial correlation (controlling for age) was run (see Figure 7). The first variable in the correlation was the behavioral advantage for congruent memories (congruent high-confidence CHR minus incongruent high-confidence CHR). The second variable was the neural activity associated to the congruent condition (Congruent ERP minus Incongruent ERP) measured at the central and parietal electrodes (C3, Cz, C4, CP5, CP1, CP2, CP6, P3, Pz, P4) that were found in the ERP cluster associated to congruence (Congruent ERP minus Incongruent ERP), from 320ms to 1176 ms. Across all subjects, a significant correlation was revealed (r = .34, *p* = .017): the greater the difference between the ERP in the congruent condition versus the incongruent condition across the central and parietal electrodes, the greater the difference between congruent high-confidence CHR and incongruent high-confidence CHR, across the participants from both age groups. Post hoc analysis revealed no statistically significant difference when we compared the correlations for the independent samples young vs. elderly subjects (p=0.47; see Figure 7). When we tested for the same correlation across both groups at the fronto-central electrodes, it did not reach significance level (*p* = 0.091). Finally, there was no correlation for the TF data.

**Figure 7.**
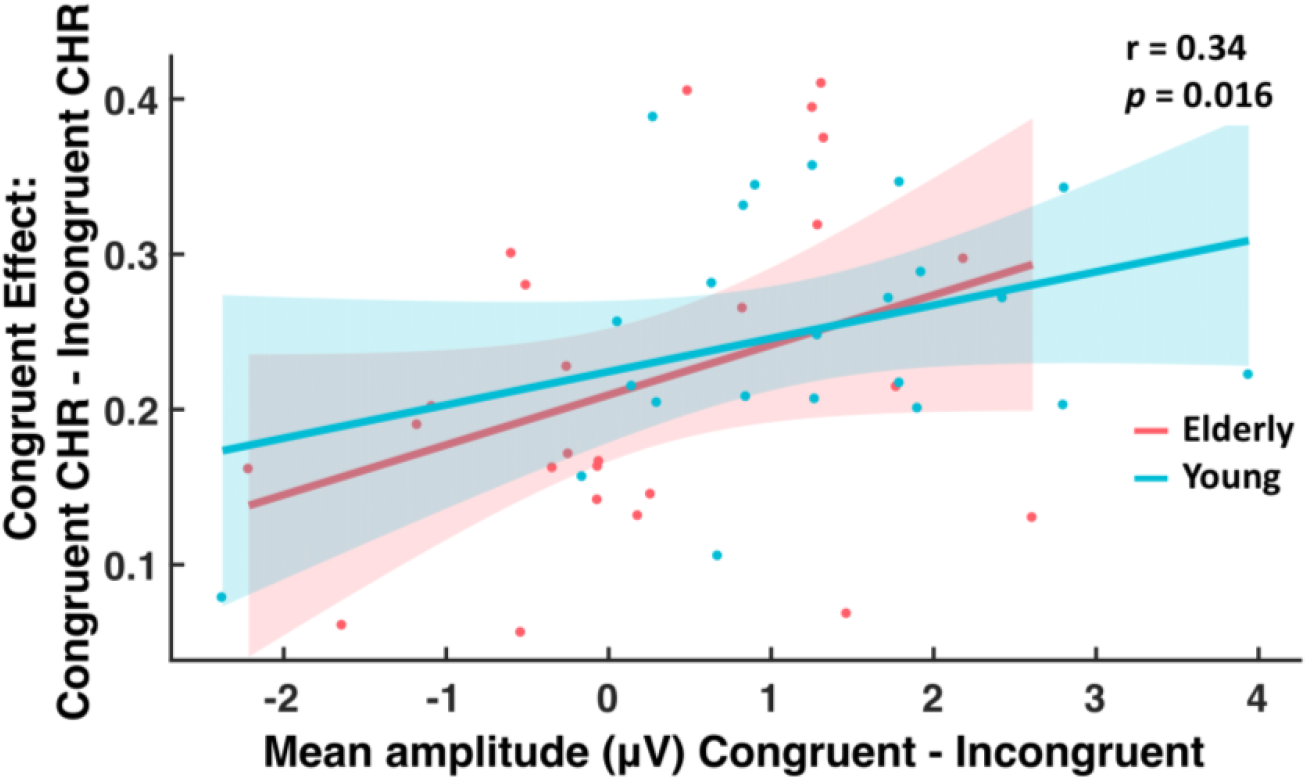
Relationship between congruence effect and ERPs. Inter-individual correlation between the ERP difference wave (mean amplitude in μV) of the congruent condition minus the incongruent condition, averaged across time (320-1176 ms after stimulus onset) and the central and parietal electrodes of the cluster (C3, Cz, C4, CP5, CP1, CP2, CP6, P3, Pz, P4), with the difference resulting from congruent high-confidence CHR minus incongruent high-confidence CHR, across participants from both age groups controlling for age. The lines show the fit of a generalized linear model to the data for each age group separately, with the 95% confidence bounds displayed. The ERP congruent minus incongruent difference wave predicted the difference in congruent memory performance across individuals in both groups controlling for age (Partial correlation; r =.34, p=.016).

## 3. Discussion

We investigated how age impacts on the semantic congruence effect and its relationship to event-related potentials (ERPs) as well as neural oscillations. Our results demonstrate that semantic congruence boosts longterm memory in both age groups, but this effect is less pronounced in the elderly. At the neural level, this observation was paralleled by age-related differences in post-stimulus neural activity, including large differences in ERP amplitude and differences in the relative power of brain oscillations in the theta-alpha and low beta range. Importantly, ERP differences associated with congruent semantic matches across central and parietal electrodes (320-1176 ms after stimulus onset) predicted the increases in memory performance for congruent items across participants from both age groups (i.e., people with greater congruency ERPs remember congruent information better, see Figure 7). Thus, our results suggest that the semantic congruence effect declines with age and they provide evidence for a role of ERPs, theta-alpha and low beta oscillations as underlying neural mechanisms.

The elderly still benefit from semantic congruence during encoding, but this effect is reduced in comparison to younger subjects (Figure 1C). This observation was expected and is largely compatible with previous findings investigating the effect of semantic processing on memory performance in young and older adults (Daselaar et al., 2003; Erber, 1974; Eysenck, 1974; Naveh-Benjamin et al., 2002; Simon, 1979). However, Crespo-Garcia et al. (2012) have used face-pair locations and found that both age groups benefitted similarly from the semantic congruence effect, but this did not compensate for the age-related reduction in overall memory (i.e. age similarly affected congruent and incongruent conditions). Yet, others suggest that age-related memory decline can be compensated for by semantic processing (Badham et al., 2012; Naveh-Benjamin et al., 2005; Patterson et al., 2009). One possible explanation for the mixed findings may relate to differences in task demands. Specifically, here the subjects engaged in a deep semantic encoding task with congruent vs incongruent word judgements, providing evidence for a reduced congruence effect in the elderly. In contrast, when semantic relatedness has been found to benefit older adults’ memory more than younger adults, subjects were not instructed to actively relate the word pairs during the encoding phase (Badham et al., 2012; Naveh-Benjamin et al., 2005; Patterson et al., 2009). Future studies may address this point more directly.

The congruence effect for high-confidence memory responses declined with age, but this interaction was only significant across both experiments and in the behavioral group alone (Figure 1). In other words, in the EEG group such age-related differences in memory performance were not apparent. This is in line with a previous EEG study which included 30 young and 28 elderly subjects (Crespo-Garcia et al., 2012), and it suggests that sample size is critical in order to detect an interaction with age (further supported by our analysis across experiment 1 and 2). Alternatively, the absence of an age interaction could relate to interindividual differences in the underlying mechanisms of encoding congruent information. Indeed, our correlation analysis shows that the congruence effect (i.e. improved memory) varies as a function of ERP congruence differences (see Figure 7). Interestingly, this correlation was significant across young and elderly subjects suggesting one underlying mechanism of congruence encoding that may continuously change with age and/or other associated factors such as learning strategies or structural brain integrity (which have not been investigated here). However, this hypothesis needs to be addressed in a longitudinal study or a design that includes a more equally distributed age range across the life-span.

The overall ERP differences between congruent vs. incongruent items (Figure 2A) may reflect the timing of memory processes and how these are modulated by the configuration of activity in memory. In other words, they might represent how semantic congruence facilitates rapid encoding. The age-related reduction in the ERPs (Figure 2B) are suggestive of an impairment of such rapid online encoding processes. Indeed, the congruence related ERP differences predicted the behavioral advantage for congruent memories (Figure 7). In agreement with a previous study that found an interaction between congruency and memory in the ERPs (Packard et al., 2016), this further supports the notion that the congruence-related ERPs reflect processes involved in the formation of long term memories.

Reaction times were faster for the accurate classifications of congruent words as compared to incongruent words (see Table 2). This effect appeared consistently across both age groups and is in line with previous studies (Neely et al., 1989). It can be explained with models assuming that top-down connections allow high-level contextual expectancies to affect perception by preparing the visual system before the stimulus even arrives, i.e., a pre-selection of possible congruent words in semantic memory (Graboi and Lisman, 2003). This suggests that incongruent stimuli are more difficult to process than congruent stimuli, which argues against the otherwise possible explanation that the congruent memory increase was due to additional effort or difficulty during encoding.

Brain oscillations provide insights into semantic processes carried out during encoding (Hanslmayr and Staudigl, 2014). Specifically, alpha and beta power decreases during congruent (vs incongruent) items (Figure 4), as in our study, may reflect a more successful encoding possibly due to their relation to semantic deep processing, which increases subsequent memory (Craik, 2002). At the neural level, such desynchronization may be associated to an increase of information processing capability within local cell assemblies (Hanslmayr et al., 2012). Beta oscillations have also been related to the maintenance of information about recent events that facilitates integrating inputs into a larger representation in memory (Morton and Polyn, 2017). More specifically, alpha-band oscillations have been posited to reflect the key process of selective access to long-term knowledge stores which allows the semantic orientation (Klimesch, 2012) that is necessary to form semantic matches. Our results suggest the elderly may have difficulties with such cognitive mechanisms and this impairs their memory performance.

Theta oscillations play a key role in the encoding and retrieval of episodic memories (Eckart and Bunzeck, 2013; Fell and Axmacher, 2011; Fuentemilla et al., 2014; Hasselmo and Stern, 2014; Herweg et al., 2016; Sans-Dublanc et al., 2017). In our study, theta power decreases were associated with processing congruent items, further suggesting a functional role of theta in the semantic congruence effect. While this is, generally speaking, in line with previous work, the direction of the effect is opposite to what has been reported recently (Crespo-Garcia et al., 2012). More precisely, Crespo-Garcia et al. (2012) could show increased theta power for semantically related face-location associations, which, similar to our findings, changed depending on age. While there are several differences between both study designs (including stimulus material and task), this opposite pattern in theta activity is compatible with the view that neural oscillations during encoding depend on perceptual and cognitive processes of the encoding task and their relation to the subsequent memory test (Hanslmayr and Staudigl, 2014).

Changes in theta-alpha and low beta oscillations, which we observed over frontal brain regions (Figure 4), might also reflect inhibitory processes. Specifically, the “schema-linked interactions between medial prefrontal and temporal regions” (SLIMM) framework (van Kesteren, 2012) suggests that the medial prefrontal cortex detects congruency with already existing neocortical information and, in this case, inhibits the medial temporal lobe which leads to more efficient cortical learning. Indeed, congruent items were associated with lower power as compared to incongruent items and this effect was reduced in the elderly (Figure 4) who, supposedly, have structurally and functionally impaired prefrontal cortices (Hedden and Gabrieli, 2004). Clearly, neural oscillations may not necessarily originate within the prefrontal cortex; therefore, future studies may address this more directly, for instance, by using combined EEG/fMRI, which would also allow to quantify structural changes in the prefrontal cortex and medial temporal lobe.

We would like to point out that, comparable to our findings, previous memory studies did not find differential effects for the theta and (low) alpha band questioning their functional dissociation during long-term memory processes (Herweg et al., 2016). While this could also represent a spill-over between neighboring frequency bands (due to the inherent limitations of TF measurements), future studies will need to further examine this open question. Along the same lines, the behavioral effects only paralleled the pattern of neural oscillations but a correlation was observed with ERPs (Figure 7). Therefore, the supposed relationship between behavior and neural oscillations is indirect.

Finally, it is difficult to specify the full memory contents, or brain networks involved in generating and retrieving these memories. However, the advantage for congruent memories in our results cannot be caused by a better memory of the category instead of the word item, because each of the categories we used was equally paired with both congruent and incongruent words. Previous studies have shown how semantic congruence increases both memories for semantically related items (Packard et al., 2016) and episodic details (Atienza et al., 2011), suggesting that semantic congruence does indeed enhance both the specific episodic and the semantic information content of memories. Future studies might help clarify this question.

Taken together, the beneficial effect of semantic congruence on long-term memory is less pronounced in the elderly. A positive ERP and a later decrease in theta-alpha and low beta power during online encoding accompanied these age-related differences suggesting a functional link. Therefore, memory impairments during healthy aging may be partly explained by changes in the neural mechanisms that underlie the rapid encoding of semantically congruent memories, which depend on neural oscillations within low to mid frequency bands.

## Acknowledgements

This work was supported by the German Research Foundation (Deutsche Forschungsgemeinschaft Grant BU 2670/7-1 to N.B.)

We are grateful to Maxi-Sophie Kuhlmey and Ramona Reineke for their help collecting data.

The authors declare no competing financial interests.

